# An integrative analysis to predict the Active Ingredients and explore Polypharmacological Mechanisms of *Orthosiphon stamineus Benth.* for human health

**DOI:** 10.1101/2021.06.12.448214

**Authors:** Xingqiang Wang, Weiqing Zhao, Xiaoyu Zhang, Zongqing Wang, Chang Han, Jiapeng Xu, Guohui Yang, Jiangyun Peng, Zhaofu Li

## Abstract

**Background:** *Orthosiphon stamineus Benth.* was a dietary supplement or a traditional folk herb with widespread clinical application, but it was lack of comprehensive understanding of its active ingredients and polypharmacological mechanisms. The aim of this work was to systematically study its natural compounds and the molecular mechanisms of the *O. stamineus* via network pharmacology.

**Methods:** Compounds from *O. stamineus* were collected by literature retrieval and evaluated by SwissADME with the physicochemical property (ADMET model) and the likelihood for a natural medicine. The connection of active ingredients and target genes was built and confirmed by Cytoscape and AutoDock vina. Then, Gene Ontology (GO) enrichment analysis and Kyoto Encyclopedia of Genes and Genomes (KEGG) pathway analysis were separately performed to obtain a more in-depth understanding of *O. stamineus*. Finally, the relationship among active ingredients, targets, and diseases was built to clarify the polypharmacological mechanisms and found relevant active substances for further drug discovery.

**Results:** A total of 159 compounds of *O. stamineus* were collected, and 22 potentially active ingredients were screened out. The Ingredient-Target Interaction Network was built with 12 flavonoids, 3 diterpenes, 3 phenols, and 4 volatile oils, and 65 targets. The Docking analysis indicated that the ingredient-target interaction network was reliable; most ligand-receptor had a strong binding affinity (lowest binding energy: −6.9 kcal/mol). After pathway analysis, 185 significant biological processes and 36 signal pathways were found, and the ingredient-target-disease network of *O. stamineus* was constructed for polypharmacological mechanisms.

**Conclusion:** Our study clarified the polypharmacological mechanisms via the relationship among active ingredients, targets, and diseases and provided better guidance for subsequent experiments and potential active ingredients for drug discovery or health promotion.

## 1 INTRODUCTION

*Orthosiphon stamineus Benth.* (family: *Lamiaceae*), also termed *Clerodendranthus spicatus (Thunb.) C.Y.Wu* in China, is mainly distributed in tropical or subtropical areas from Southeast Asia to North Australia. This plant has white and purple flowers with stamens extending out of the corolla, which looks like cat’s whiskers, so it is also called *cat’s whiskers*, *misai kucing*, *Java tea, kumis kucing*^1, 2^. *O. stamineus* have been widely used in dietary supplements, herbal medicine and ornamental purposes particularly in Asia. In Yunnan, Guangxi and other regions of China, *O. stamineus*, as a raw material for herbal tea such as “liang cha”, has become a necessary drink for barbecue and hot pot. In Thailand, the whole plant as a dietary supplement is used for decoction to eat and the fresh leaves are pounded up and put on bruised or sprained joints for the cure. According to the records of the Dai medicine, *O. stamineus,* also named “ya nu miao” (Pinyin) in Dai language, is considered a high medicinal value that was consistently applied to treat nephritis, cystitis, rheumatism, edema, hypertension, diabetes, obesity, hepatitis, and so on^3^. In addition, many pharmacopeias such as Indonesian, Malaysia, Dutch, and French, have listed the therapeutic indications: renal cleansing and tonic kidney, used for the treatment of calculus and weight loss. And it is even employed to treat tumors, urinary tract infections and coronary artery disease in some countries^4, 5^.

Although the widespread application of *O. stamineus* in the clinical practice, it is still insufficient to support its curative effect. Plenty of pre-clinical experiments demonstrated that the various extracts of *O. stamineus* exerted pharmacological properties, including anti-proliferative^6, 7^, antioxidant^8, 9^, analgesic^10^, anti-inflammatory^9, 11^, antimicrobial^12, 13^, diuretic^14^, among others. Recently, two rare clinical trials have been performed, and the results showed that the herbal combination with *O. stamineus* seems to be useful in the reduction of discomfort and prevention of antibiotic use in women with urinary tract infections^15, 16^. Despite the fact that dozens of phytochemical studies investigated the natural compounds in *O. stamineus* from all over the world and hundreds of chemical compounds such as flavones, polyphenols, and terpenoids have been detected, very little is known about active substances, their targets, interactions, and specific molecular mechanisms. Thus, it continues to be fierce challenges for us to systematically study these natural compounds and the molecular mechanisms, which benefit further drug discoveries and functional foods.

Network pharmacology, a novel analytical method for traditional Chinese medicine (TCM) that has emerged in the last decade, could integrate systems biology, disease knowledge, pharmacology, and herbal medicine to conduct a systematic analysis on human diseases under the guidance of a holistic perspective^17, 18^. It is common knowledge that the traditional herb or herbal formulae ameliorate health via multicomponent interaction with multiple targets, regulation of multiple biological processes, and synergistic effects. This view shifts the traditional paradigm of “one disease, one target, one drug” to develop a “network-target, multicomponent therapeutics” mode^19, 20^. Fortunately, not only network pharmacology but the polypharmacology, which supports that the design or use of pharmaceutical agents that act on multiple targets or disease pathways, also coincides with traditional medicine^21^. Therefore, network pharmacology is a very appropriate technology to explore herbs of Dai medicine systematically.

In view of the shortcomings of the current researches of *O. stamineus*, in our study, we scrutinized related studies and collected almost all compounds of *O. stamineus.* Then, we evaluated the physicochemical property, oral bioavailability, and drug-likeness for the accumulation of phytochemical information by network pharmacology tools. Finally, the relationship among active ingredients, targets, and diseases was built to clarify the polypharmacological mechanisms and found relevant active substances for further drug discovery or health promotion.

## 2 MATERIAL AND METHODS

The workflow of this study was depicted in Figure 1.

**FIGURE 1.**
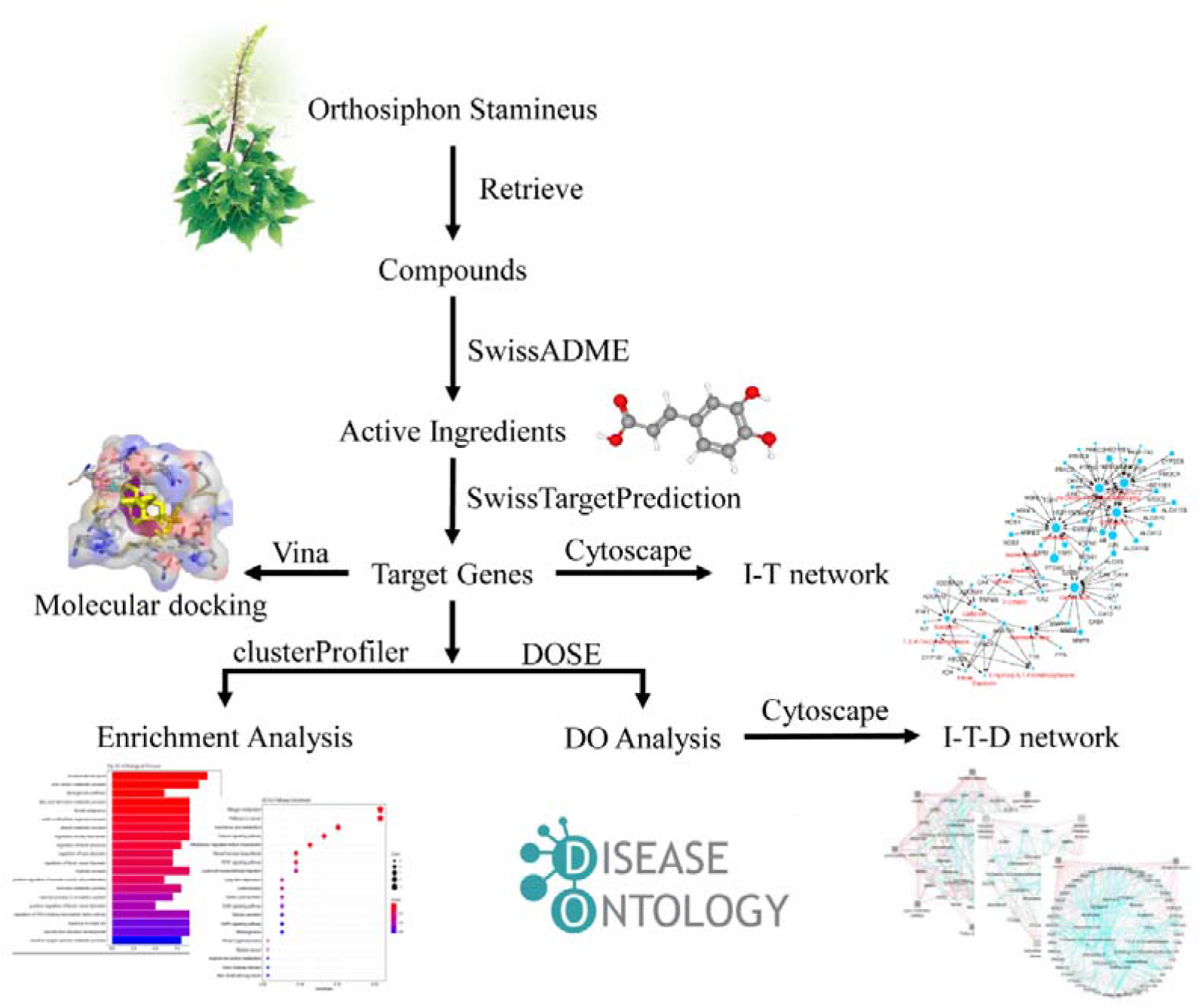
Diagram of methodology applied in this study.

### 2.1 Compounds Database Building and ADMET Evaluation

In order to collect the information of compounds from various extracts of *O. stamineus*, we retrieved literature in PubMed, Embase, Google Scholar, and Chinese databases, including CNKI (http://new.oversea.cnki.net/), WANFANG data (http://www.wanfangdata.com.cn/), and SinoMed (http://www.sinomed.ac.cn/). The keywords were including “Orthosiphon stamineus,” “clerodendranthus spicatus,” “Java tea,” “ingredient,” and “compound.” Then got the compound CID, SMILES identifier of compounds collected via PubChem (https://pubchem.ncbi.nlm.nih.gov/) and constructed compounds database of *O. stamineus* by Microsoft Excel 2013. ADMET is an acronym in pharmacology for absorption, distribution, metabolism, excretion, and toxicity, and it plays an indispensable role in drug discovery. Here, we conduct a web tool SwissADME (http://www.swissadme.ch/) to evaluate ADMET parameters and drug-likeness^22^.

### 2.2 Screening Potential Active Ingredients

SwissADME provided various chemical properties of small molecules, including water solubility, gastrointestinal absorption, drug-likeness (DL), and oral bioavailability (OB)^22, 23^. *O. stamineus* was usually made as a decoction to intake, so we screened the potential active ingredients via the following indicators: (i) Log S > −6 or Solubility class was at least moderately soluble, (ii) the compound must fully meet Lipinski‘s Rule of Five^24^, (iii) and OB ≥ 30%.

### 2.3 Target Genes and Protein-Protein Interaction Network Prediction

Based on the SMILES identifier, target genes of each potential active ingredient were predicted by SwissTargetPrediction (http://www.swisstargetprediction.ch/) with “Homo sapiens” setting^25^. The SwissTargetPrediction calculated the probability in the prediction output page from the combined score of the compound most similar to the query molecule with a given protein as a target^26^. The greater the probability is, the more accurate the predicted target is. We set a probability of 0.2 as the threshold to filter more credible target genes.

To obtain protein-protein interaction networks, we inputted the filtered target genes into STRING (https://string-db.org/) with “Homo sapiens” as the “organism,” and protein-protein interaction network was obtained by setting “minimum required interaction score” as “high confidence (0.700)” during conducting STRING^27^.

### 2.4 Gene Function and Pathway Enrichment Analysis

In order to better understand the biological effects of active ingredients to target genes, Gene Ontology (GO) enrichment analysis and Kyoto Encyclopedia of Genes and Genomes (KEGG) pathway analysis were separately performed using the R package “clusterProfiler”^28^. The qvalue was the corrected P-value by Benjamini-Hochberg (BH) method, and it was significant for a qvalue less than 0.05.

### 2.5 Disease Ontology Analysis

The Disease Ontology (DO) is a standardized ontology of human disease for the examination and comparison of genetic variation, phenotype, protein, drug, and epitope data (https://disease-ontology.org/)^29^. The DO analysis aims to analyze which human diseases (or class of diseases) are enriched in my gene or protein sets by hypergeometric tests. In terms of target genes above, we performed the DO analysis via the R package “DOSE”^30^. The qvalue was the corrected P-value by Benjamini-Hochberg (BH) method, and it was significant for a qvalue less than 0.05.

### 2.6 Network Construction

The ingredient-target-disease (I-T-D) network was established via the Cytoscape 3.7.1 through three steps as follows^31^: (i) the active ingredient-target network was generated by linking the potential active ingredients, target genes, and protein-protein interaction; (ii) the target-disease network was constructed by extracting the results of DO analysis; (iii) the effects or relationships from active ingredient to disease were analyzed by segmenting the I-T-D network into subnetworks based on disease classification. In these results, nodes stand for active ingredients, target genes or diseases, and edges represent these three interactions. So the degree value of the node is the edge numbers of ingredients or genes in the network, and it represents the importance of the ingredients or genes.

### 2.7 Molecular Docking Analysis

To validate the reliability of the ingredient-target associations, we performed molecular docking analysis using AutoDock vina 1.1.2 (http://vina.scripps.edu/index.html)^32^. Here, we only take some hub target genes and ingredients for example. Firstly, ingredients (ligands) were downloaded from the PubChem database with SDF format, and the target proteins (receptors) were collected from the RCSB website (http://www.rcsb.org/) in PDB format. Then the SDF files of ligands were converted into PDB format, and the original crystal ligands and water molecules were removed from receptors. As preparation for docking, both the receptors and the ligands were added hydrogens, computed Gasteiger charges, and the non-polar hydrogens were merged. The docking site was defined within a grid box around residues of the binding site, and the grid space was set as 1 Å. Finally, the AutoDock vina 1.1.2 was performed to explore the conformational binding site and compute binding energies using the Lamarckian genetic algorithm, and the other parameters (e.g., the random seed was set as 708076952 and the number of iterations was set to100 times) were the same for each docking.

### 2.8 Statistical Software

The statistical analysis was performed using R 3.5.0 (2018/04/23), Bioconductor 3.7, MGLTools 1.5.6 (http://mgltools.scripps.edu/), AutoDock vina 1.1.2, Pymol v2.3.0, and Cytoscape 3. 7.1 on Windows System x64.

## 3 RESULTS

### 3.1 Potential Active Ingredients

Through literature retrieval and evaluation, we collected 159 compounds, including terpenoid, flavonoid, aromatics, phenol, alkyls, organic acid, and others. These compounds were all listed in Supplementary table S1. After the calculating in web tool SwissADME, we screened out 62 potentially active ingredients (listed in Supplementary table S2) applying to the screen criteria above. Particularly, volatile oils account for 35 of the 62 active ingredients in the *O. stamineus* (about 56.45%), terpenoids account for 21 (33.87%), flavonoids account for 12 (19.35%), indicating that these three chemicals were the main active ingredients in *O. stamineus*.

### 3.2 the ADME Parameters and Physicochemical Properties of These Ingredients

As shown in Supplementary table S2, the average molecular weight of these ingredients was about 230.59, the median was 185.22, the maximum was 492.56, and the minimum was 100.16. The prediction results of drug characteristics shown that of the 62 active ingredients, 77.42% had good water solubility, 96.77% had good intestinal absorption, and 74.19% can permeate the blood-brain barrier (BBB). There were 29 small molecules predicted as non-inhibitor of five major cytochromes P450 (CYP) isoforms. All the 62 active ingredients had high OB value; for example, the OB value of 59 compounds was 55%, and the three compounds were 56%. All the 62 compounds complied fully with “Lipinski’s rule of five,” indicating that these had great drug-likeness. These results demonstrated that *O. stamineus* was reasonable as a decoction to intake.

### 3.3 Ingredient-Target Interaction Network

In order to understand Ingredient-target (I-T) interaction mechanisms in the human body, we connected potential active ingredients with target proteins or genes via conducting SwissTargetPrediction and STRING. In the results, 65 target genes (proteins) were linked to 22 active ingredients, as shown in Supplementary Table S3 and Table S4. We used 22 potentially active constituents and relevant targets to generate a bipartite graph that illustrating the interaction between compounds and their target proteins, target proteins, and target proteins (Figure 2). The I-T network contains 87 nodes and 238 edges, including 12 flavonoids, 3 diterpenes, 3 phenols, and 4 volatile oils. In this I-T network, 14-Deoxo-14-O-acetylorthosiphol Y, Orthosiphol Y, Orthosiphol Z, Vomifoliol, and Caffeic acid were all surrounded with 15 target proteins and were identified as the center of the network to suggested which importance in pharmaceutical effect. Likewise, some targets that shared multiple constituents were considered as hub target proteins, such as ABCG2 and AKR1B1 were a target of 11 compounds, respectively. OPRD1 also was the target of 8 flavonoids. Additionally, 4 essential oils only map to 4 targets depicted in Figure 2, so It was insignificant for O. stamineus to cure disease, due to which mainly were present in fresh plants and were easily volatile during processing.

**FIGURE 2.**
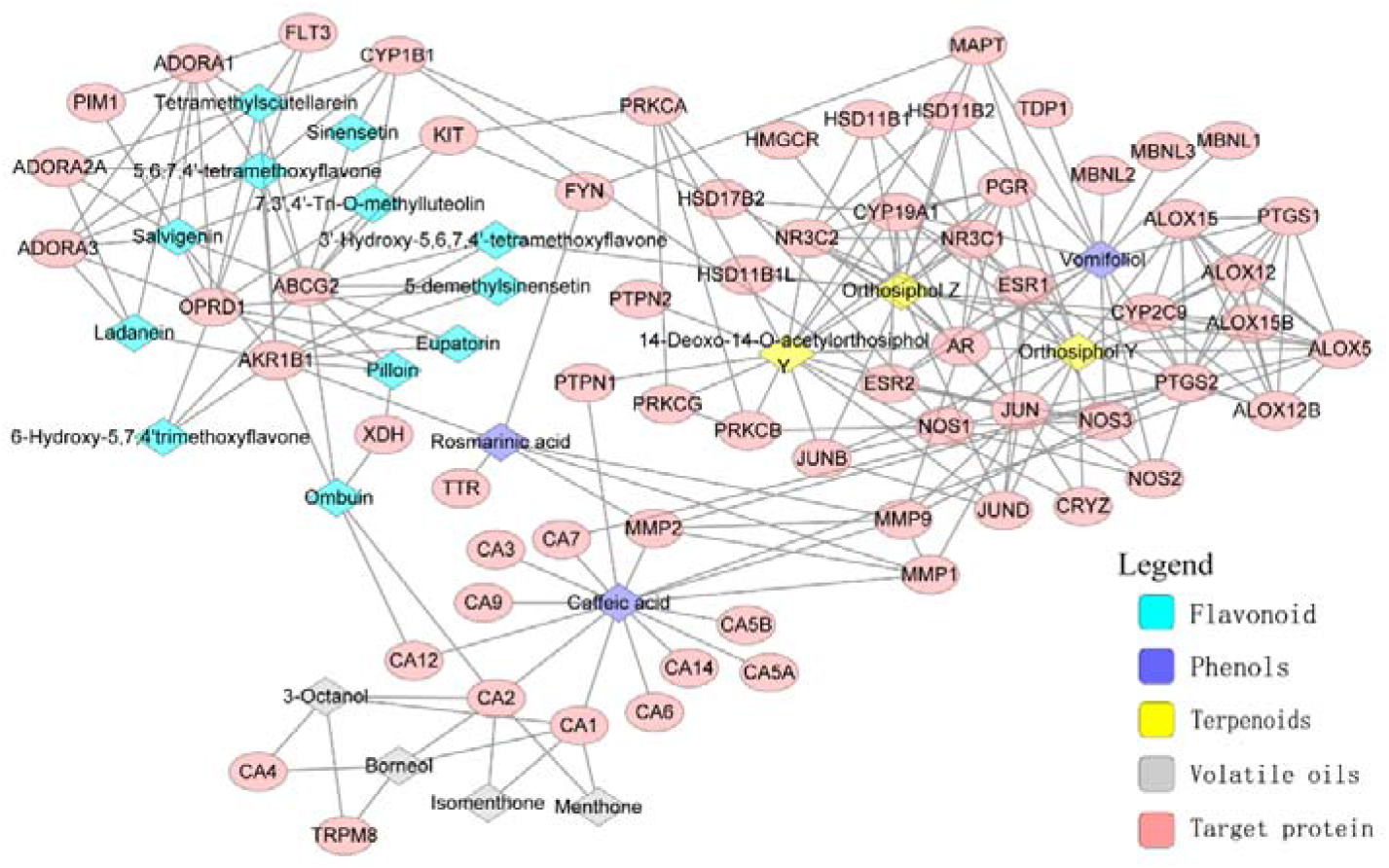
The ingredient-target interaction network of *O. stamineus.* The pink ellipses represent target genes, and the colorful diamonds stand for potential active constituents in *O. stamineus*.

### 3.4 Molecular Docking Analysis

Forty-five pairs of the ligand-receptor complex were analyzed to access the relative binding affinity of the ligand toward the receptor and predict the binding sites in our silico study. Based on the docking analysis results shown in Table S5, the binding free energies ranged from −6.9 kcal/mol to −16.3 kcal/mol. The hub target AKR1B1 had 11 ligands such as 6-Hydroxy-5,7,4’trimethoxyflavone, Salvigenin, Eupatorin, Pilloin, Ladanein, Rosmarinic acid, and their binding energies were −10.9,−11.3, −11.5, −12.1, −11.9, and −13.2 kcal/mol, respectively. Particularly, Orthosiphol Z may bind to many targets such as AR, NR3C1, NR3C2, PGR, and it all has the highest binding affinity (docking score: −14.5 to −16.3, Figure 3). In brief, these docking simulations indicated that the ingredient-target interaction network was of reliability, and most ligand-receptor had a strong binding affinity, especially Orthosiphol Z.

**FIGURE 3.**
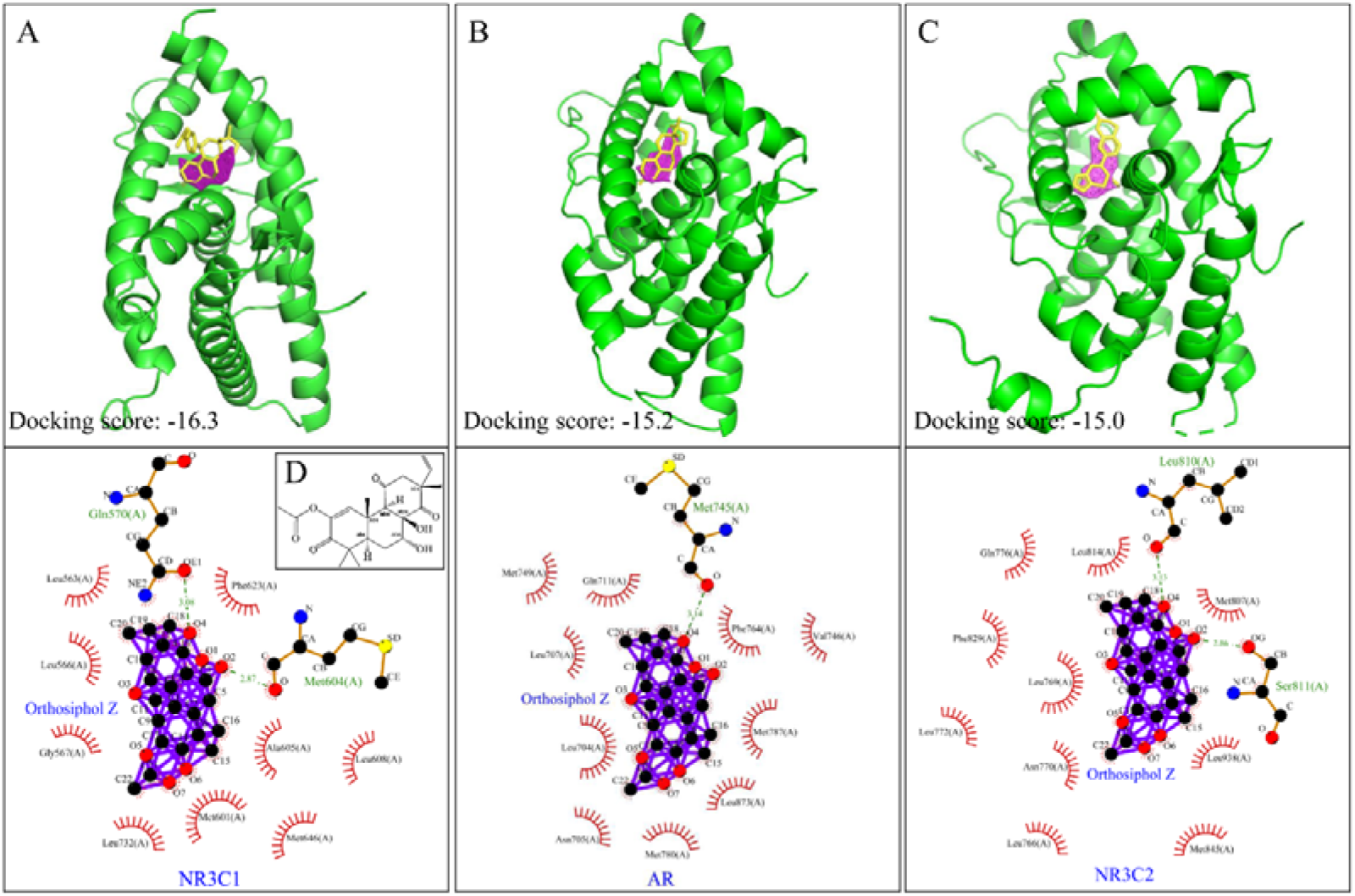
The top 3 ligand-protein interaction diagrams of the structural interactions of active ingredients (purple) and target protein with their inhibitors (yellow). The upper panel shows the 3D structure of the orthosiphol Z (purple) and target protein (green cartoon) with their inhibitors (yellow); the inhibitors and targets are (A) mifepristone and NR3C1, (B) metribolone and AR, and (C) LD1 and NR3C2, respectively. In the bottom diagrams depicted by ligplot+, the dashed lines indicate hydrogen bonds; the spoked arcs in red indicate hydrophobic contacts. Carbons are in black, nitrogens in blue, oxygens in red and orthosiphol Z in purple. Here, (A) The interactional model of NR3C1 and orthosiphol Z, (B) AR and orthosiphol Z, (C) NR3C2 and orthosiphol Z were shown. (D) The chemical structure of orthosiphol Z.

### 3.5 Potential Mechanism

In this section, we input 65 target genes into the “clusterProfiler” for the biological effects of 22 active ingredients and presented the results of GO enrichment analysis and KEGG pathway enrichment analysis in Supplementary Table S6 and Table S7. In these results, one hundred eighty-five significant biological processes and 36 signal pathways were found out, and we selected the top 20 terms (ranked via adjusted P-value) for charting. As figure 4A illustrated, the potential drug targets ware major related to human biological process including “bicarbonate transport,” “arachidonic acid metabolic process,” “long-chain fatty acid biosynthetic process,” “fatty acid derivative metabolic process,” “hormone metabolic,” “regulation of blood pressure,” “reactive oxygen species metabolic process” among others. The 36 signal pathways listed in Table S6, such as “Nitrogen metabolism,” “Arachidonic acid metabolism,” “Aldosterone-regulated sodium reabsorption,” “VEGF signaling pathway,” could be related to the potential polypharmacological mechanism of *O. stamineus* (Figure 4B).

**FIGURE 4.**
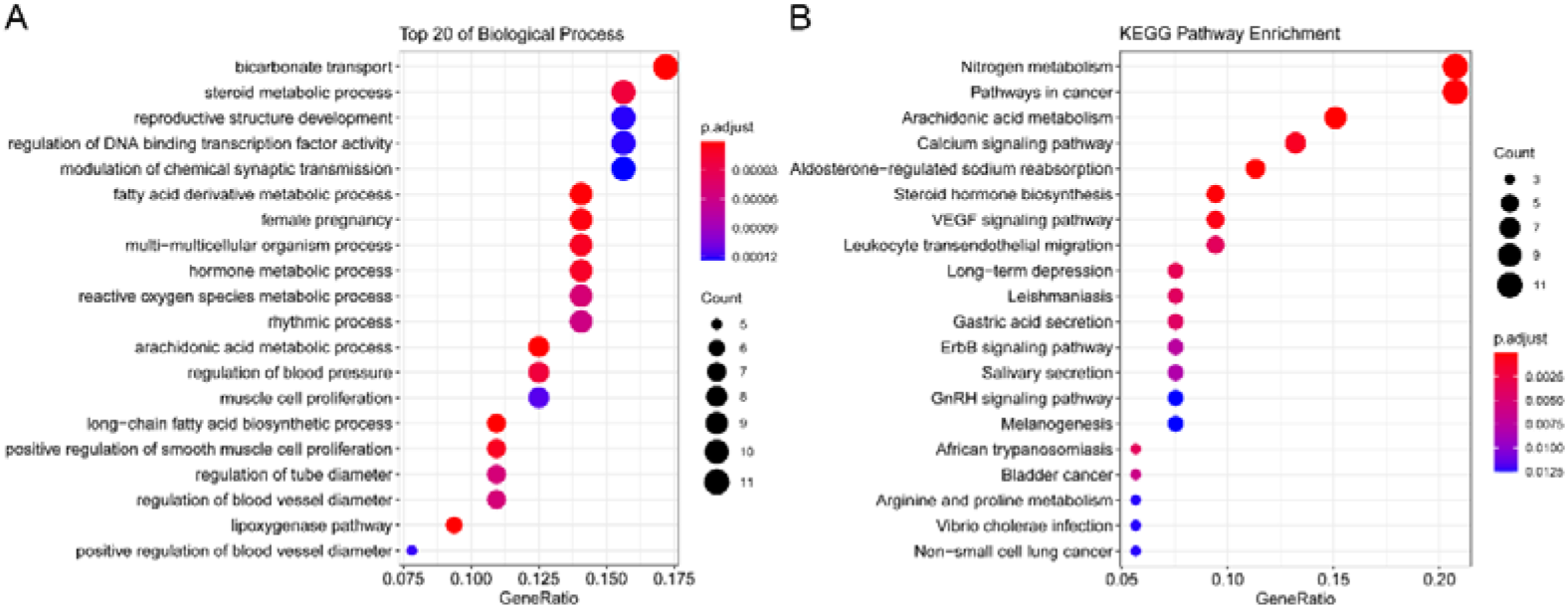
The dot plot of enrichment with target genes. (A) GO enrichment analysis (top 20) of the target proteins (*P* < 0.05). (B) KEGG pathways of target proteins (*P* < 0.05).

### 3.6 Unraveling the Molecular Basis of Diseases Treated by *O. stamineus*

*O. stamineus*, a valued traditional folk medicine, has been widely used for several human disease or conditions such as edema, hypertension, nephritis, diabetes, hyperlipidemia, urinary inflammation, calculus, gout, and rheumatism, because of their diuretic, antioxidative, anti-inflammatory, analgesic, cytotoxic, antihypertensive, hypolipidemic property. Here, we connected the drug-target network with diseases by DO analysis since it could be an effective way to comprehensively understand the *O. stamineus.* Assuming that drug components acting on the same protein in the same network can be linked to multiple diseases and treat these diseases finally, this disease-centric network may provide more evidence and novel ideas on applying for traditional medicine.

#### 3.6.1 Regulating Metabolism

The annotation by DO analysis shown that of the 22 active ingredients, 16 linked to some nutritional and metabolic diseases, which main contained type 2 diabetes mellitus, lipid metabolism disorders (familial hyperlipidemia, hyperglycemia, and Fabry disease), and nutritional disease (obesity, overnutrition). In the light of Figure 5, there are 21 targets related to the nutritional and metabolic diseases, and the hub target proteins NR3C1, AKR1B1, PGR, ALOX12, PTGS2, and NOS3 had the most degree in this network. They were connected to lipid metabolic disorders and obesity tightly. It depicted the *O. stamineus* could treat these metabolic diseases through regulating these targets (Figure 5A).

**FIGURE 5.**
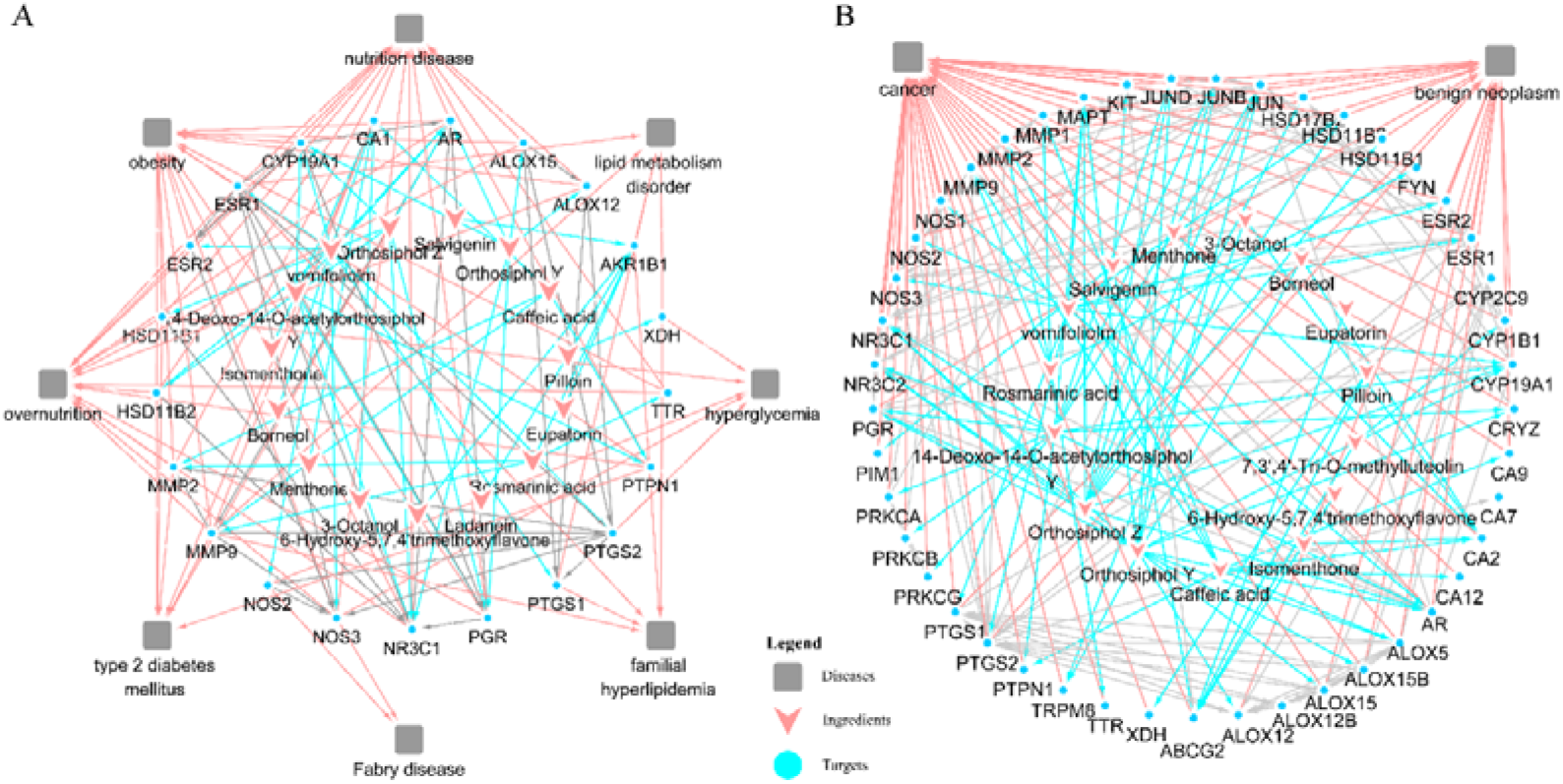
The Ingredient-target-disease (I-T-D) network of *O. stamineus*. (A) The connection of among active compounds, their targets, and metabolic diseases or (B) neoplasm.

#### 3.6.2 The Molecular Basis of Antiproliferative

O. stamineus was rarely used for neoplasm in the history of folk medicine. However, our results showed that 17 active ingredients and 42 target genes concerned with benign neoplasms and cancers (figure 4B). Figure 5 illustrated that Orthosiphol Y, Orthosiphol Z, and 14-Deoxo-14-O-acetylorthosiphol Y were the core compounds, and the hub genes such as ALOX12, JUN, JUND, PTGS2, PGR, ABCG2, NR3C1 were all the critical molecules for antiproliferative. Thus, through regulating of these targets, O. stamineus could regulate some signals (i.e., nitrogen metabolism, pathways in cancer, non-small cell lung cancer, VEGF signaling pathway, ErbB signaling pathway, Wnt signaling pathway, and others) for antineoplastic effects (the detail information was listed in Table 1).

**Table 1.**
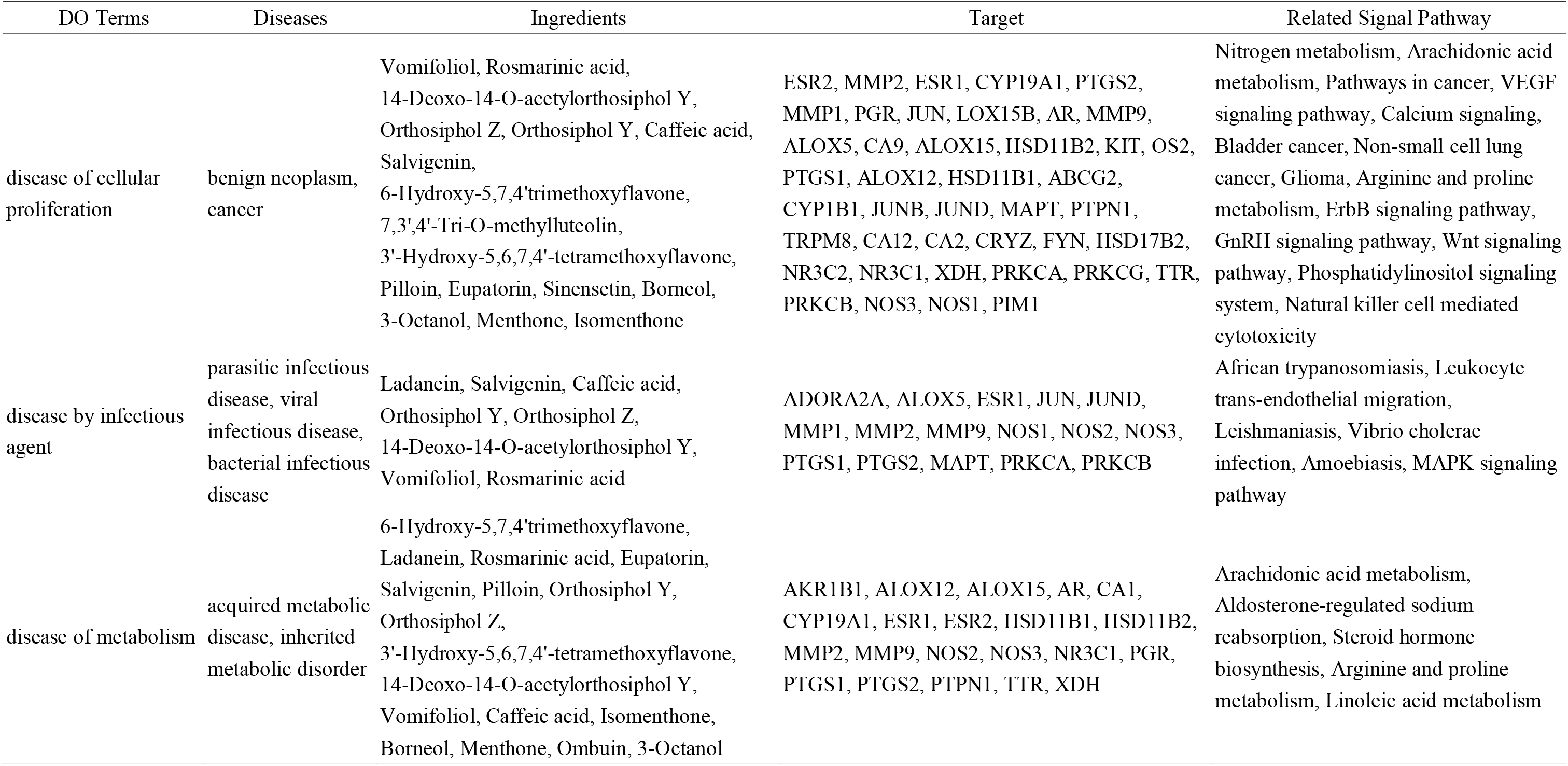
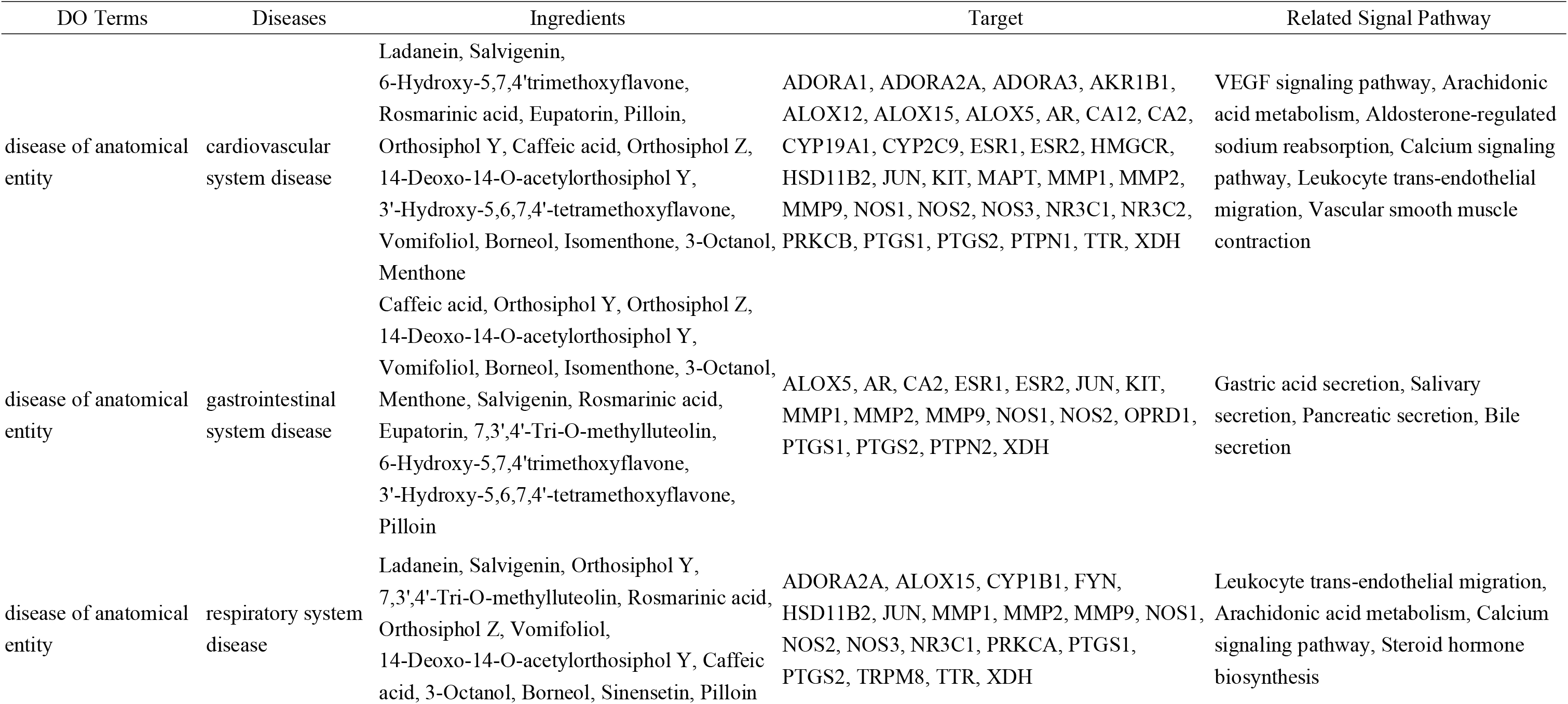
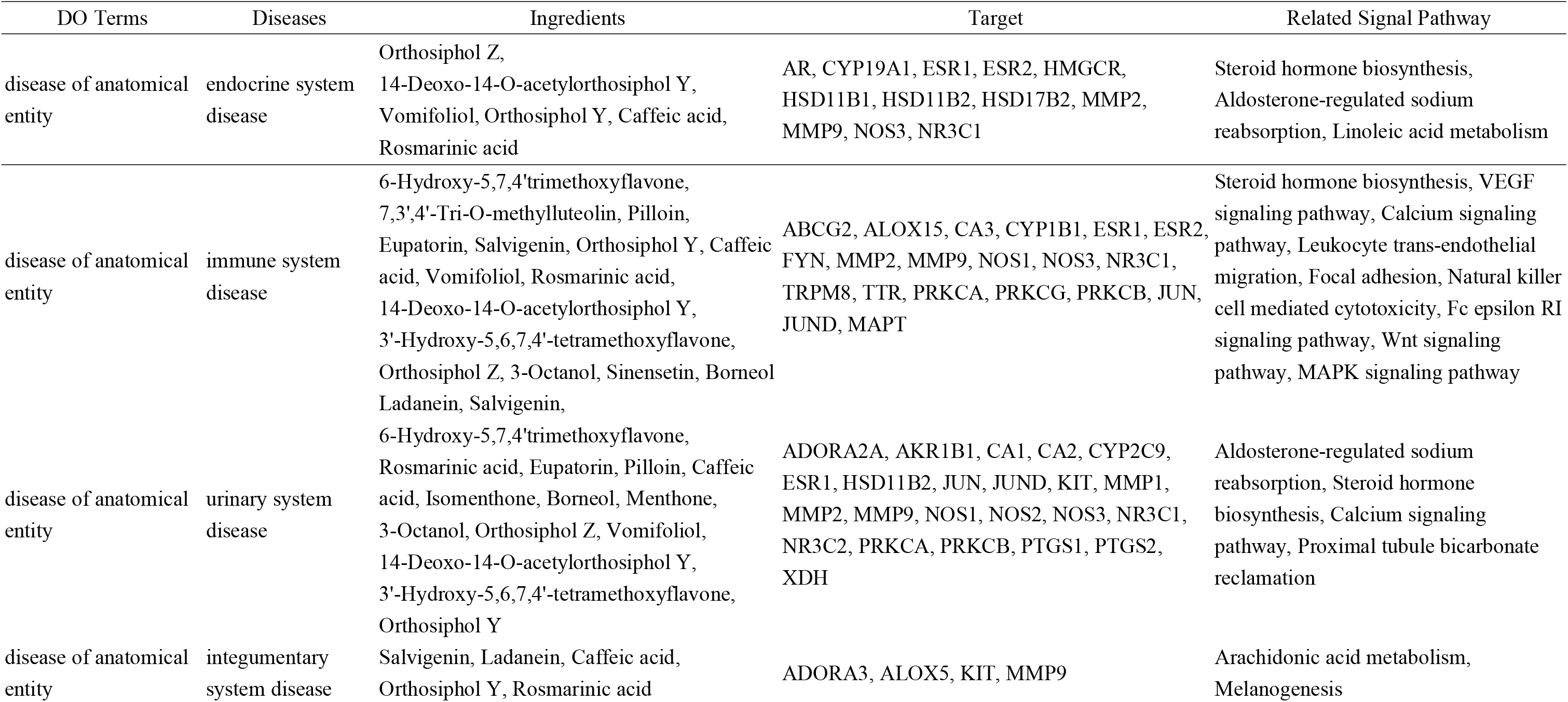
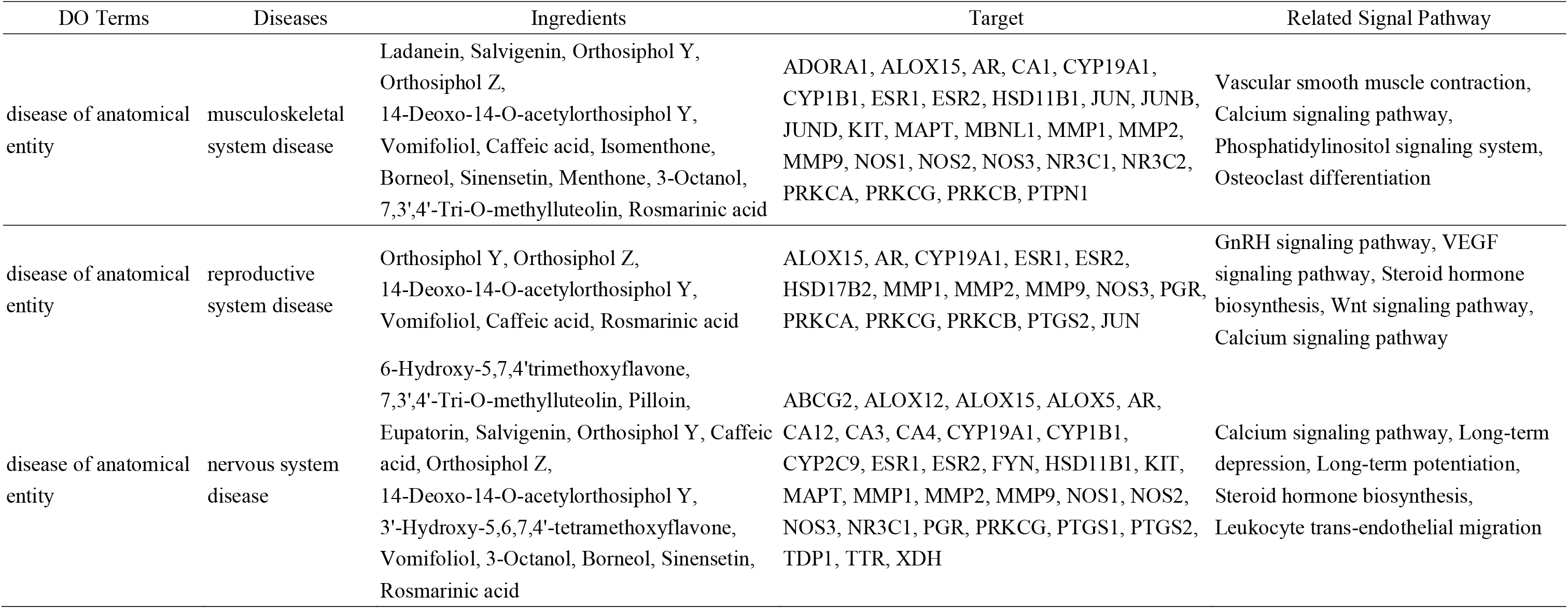
The potential active ingredients from *O. stamineus* and their targets, signal pathway and treated diseases

#### 3.6.3 Anti-inflammatory and Anti-Infectious Property

As can be seen from Table 1, 18 target genes and eight constituents had an association with infectious diseases, which suggested that these molecules could play a vital role in treating infection or inflammation. Consideration with the resultant of pathway analysis such as “Leukocyte transendothelial migration,” “MAPK signaling pathway,” “African trypanosomiasis” above, we believe that eight compounds Ladanein, Salvigenin, Caffeic acid, Orthosiphol Y, Orthosiphol Z, Vomifoliol, Rosmarinic acid, and 14-Deoxo-14-O-acetylorthosiphol Y could regulate essential proteins (i.e., MAPT, PRKCA, PRKCA, PRKCB, JUN, JUND, NOS2, PTGS2, etc.) to promote our body’s resistance against the infectious agent and to remain balance of immunity.

#### 3.6.4 Other Conditions

Besides the results described above, we have listed ten diseases classified by anatomy based on the anatomical structure in the DO database. These ten diseases were mapped to 22 potentially active ingredients and 65 target genes totally in our study, and the details could be found in table 1.

Some essential indications in table 1 were shown that O. stamineus could treat cardiovascular disease, gastrointestinal disease, respiratory disease, endocrine disease, urinary system disease, immune system disease, nervous system disease, and so on through binding related targets, and some of them shared same targets or same signaling pathways. For instance, MMP9 was shared by almost all categories as well as PTGS, NOS, ESR, NR3C1, JUN, KIT, ALOX15 have a higher frequency in these ten diseases (Table S9); likewise, several signaling pathways, such as Arachidonic acid metabolism, VEGF signaling pathway, Calcium signaling, Leukocyte trans-endothelial migration, Aldosterone-regulated sodium reabsorption, Steroid hormone biosynthesis, were likely to be involved in most disorders from our DO analysis.

## 4 DISCUSSION AND CONCLUSION

*O. stamineus* was extensively used for edema, hypertension, diabetes, obesity, rheumatism, nephrolithiasis, urinary inflammation, hyperlipidemia by decocting or making herb tea in our traditional folk medicine^3, 4^. So far, accumulating evidence indicates that it has several pharmacological functions, including diuretic, antioxidant, anti-inflammatory, antimicrobial, anti-proliferative, anti-angiogenesis, hepatoprotective activities and ameliorate metabolism^10^. In this study, we collected 139 compounds and evaluated its by ADMET model, then predicted water solubility, gastrointestinal absorption, blood-brain barrier permeation, drug-likeness (DL) and oral bioavailability (OB), screened potential ingredients and conducted their targets prediction. In the end, we generated several ingredient-target-disease networks to explore potential pharmacological mechanism and extend understanding of *O. aristatus* for drug discovery.

Consideration for good water solubility, high oral availability, and excellent drug-likeness, we screened out 62 potentially active ingredients, including 35 volatile oils (56.45%), 21 terpenoids (33.87%), 12 flavonoids (19.35%), and several phenols. In general, volatile oils consist of terpenoids, aliphatic, and aromatic compounds and their oxygen-containing derivatives and could sustain loss during boiling and drying^33^. Moreover, flavonoids, terpenoids, and phenols all have anti-inflammatory, analgesic, anti-microbial, antiproliferative activities, and immune regulation, as do volatile oils. Thus, it indicated that the main active ingredients of *O. aristatus* were volatile oils, terpenoids, flavonoids, and several phenols and suggested us optimizing the processing technology and the usage for avoidance loss of active ingredients.

Results of targets prediction and molecular docking demonstrated that the therapeutic effects of *O. aristatus* were in association with 22 ingredients, and the relationship of ingredient-target interaction was great reliable because most ligand-receptor had a strong binding affinity. Based on the binding free energies of docking, the ingredients Orthosiphol Z, Orthosiphol Y, and Rosmarinic acid all have excellent binding affinity to their target proteins (Table S5, Figure 3). Till to date, there is not any published paper about the pharmacological activity of orthosiphol Y, Z, and only the orthosiphol A, an isomer of orthosiphol Y, Z, has been proven to benefit diabetes and gout due to that retard intestinal maltase function^34^ and promote UA excretion^10^. However, the water solubility of the Orthosiphol A is poor, and the gastrointestinal absorption is low in our ADMET model. Different from Orthosiphol A, Rosmarinic acid, a water-soluble and absorbable polyphenol that been researched endlessly, had several exciting effects, including antioxidant^35^, anti-inflammatory^36^, anti-tumor^37^, antimicrobial^38^, anti-diabetic^39^, neuroprotective^40, 41^ and nephroprotective activities^42,43^ even cosmetic effects^44^. Similarly, two flavonoids Eupatorin and Sinensetin also been confirmed to possess almost the same pharmacological effects as *O. aristatus*^4, 45–47^. Noteworthily, our results have shown that Rosmarinic acid cannot cross the blood-brain barrier and its gastrointestinal absorption was low (Table S2). But the pre-clinical experiments showed that it can improve central system diseases such as epilepsy and Alzheimer’s disease^40, 41, 47^, which makes me confused. Anyway, we speculate that the O. aristatus own many ponderable ingredients for phytochemical screening and validation.

In view of the pharmacological mechanism of *O. aristatus*, we focus on the elucidation of the anti-neoplastic and metabolism-regulative effects for more valuable opinions. In past decades, accumulated researches have evaluated the antineoplastic effects of *O. aristatus* in vivo and in vitro, and yet *O. aristatus* rarely used for neoplasm in traditional medicine. these researches showed that the standardized extracts of *O. aristatus* could inhibit the growth of hepatocellular carcinoma, fibrosarcoma, and colorectal carcinoma cell line by inducing apoptosis^6, 7, 48^ or antimutagenic efficacies^49^. The chloroform extract of *O. aristatus* had solid inhibitory activities against five cancer cell lines, and Luo *et al*.^50^ concluded that the backbones of diterpenoids had a significant influence on cell toxicity due to the six active compounds (Spicatusene B, Spicatusene C, Orthosiphol R, Orthosiphol N, Orthoarisin A, Orthosiphonone D). Ahamed *et al*.^7^ further studied the antiproliferative activities of the ethanolic extract of O. aristatus in athymic mice with colorectal tumor, and it caused 47.62 ± 6.4% and 83.39 ± 4.1% tumor growth suppression after four weeks at the dose of 100 and 200 mg/kg body weight, respectively. Interestingly, the ethanolic extract of *O. aristatus* could significantly reduce the vascular endothelial growth factor (VEGF) level and suppress the tumorous angiogenesis via the VEGF-VEGFR signal^7^, which has been depicted in our results (Figure 4, Table S7). Another research suggested that the methanolic extract of *O. aristatus* synergistically enhanced cytotoxic effect towards MCF-7 hormone-sensitive human breast cancer cells with Tamoxifen. Still, the extract itself did not show any anticancer efficacy^51^. Here, the synergistic effect of *O. aristatus* was associated with the hormone metabolic process as mentioned above. In Table 1, We have listed some abnormal signal pathways in tumors from our I-T-D network for a comprehensive understanding of the potential anti-tumor mechanisms of O. aristatus. Furthermore, based on our I-T-D network analysis of O. aristatus, those candidate signaling such as “Nitrogen metabolism,” “Pathways in cancer,” “Arachidonic acid metabolism,” “VEGF signaling pathway,” “Non-small cell lung cancer,” “ErbB signaling pathway,” “Wnt signaling pathway,” and even “the Calcium signaling,” will provide a basis for further pharmacological verifications with confirmation of active ingredients.

In addition, concerning the effect of *O. aristatus* on metabolic diseases, three studies^52–54^ indicated that the aqueous extract could ameliorate blood glucose level and lipid profile in diabetic rats as well as did work in pregnant diabetic rats. They further investigated the primary mechanisms that indicated *O. aristatus* probably increases the sensitivity of pancreatic beta cells to insulin or (and) peripheral utilization of glucose by the tissues for lowering plasma glucose^52, 55^. Meanwhile, Chen and Xu *et al*.^10, 56^ evaluated the function of anti-gout in mice. They showed that ethanolic and ethyl acetate fraction of *O. aristatus* possessed lowering serum uric acid (UA) and relieving pain because of the increasing excretion and the suppressing production of the UA, anti-inflammation, and analgesia. These studies concluded that some active compounds in *O. aristatus* inhibited the xanthine oxidase activity and targeted proximal tubule epithelial cells for regulation reabsorption, and we did find in our research. Amazingly, an investigation demonstrated the *O. aristatus* might benefit the metabolic diseases mainly through regulating the tricarboxylic acid cycle, gluconeogenesis, lipid, and amino acid metabolism by (1)H NMR spectroscopy^57^.

Furthermore, this benefit of *O. aristatus* to our metabolism has been turned out to be reliable by a randomized clinical trial^58^. Our research also manifested a tight connection between metabolism disorders and several good water-soluble ingredients (Figure 5A, Table 1) and suggested that some pharmacological mechanisms were involved. For example, Arachidonic acid metabolism, a long-established lipid metabolism process, plays an extensive pathophysiological role in human health and diseases, aiding in cell proliferation and tissue homeostasis and in inflammatory diseases, cancer, cardiovascular diseases, and metabolism aberration, such as diabetes mellitus, obesity, lipid alteration^59^. So, it is reasonable to conduct a targeted therapy if the Arachidonic acid metabolism has been impacted by *O. aristatus*.

Besides, it seems compelling for *O. aristatus* to treat nephritis^42, 50, 56^, calculus^60^, urinary tract infection, hypertension^61^, liver damage^62^, atherosclerosis^5^, and gastritis^63^ in numerous animal experiments. A few investigations also evaluated its neuroprotection in epilepsy, depression, and Alzheimer’s disease^47^. However, most of these studies were at the pre-clinical phase; only three clinical trials of an herbal combination containing *O. aristatus* proved effective in treating hypertension, metabolic syndrome^58^, and urinary tract infections^15, 16^. Based on the I-T-D network and DO analysis in our study, it seems to give an insight that O. aristatus was a versatile therapeutic agent. Although this view is a bit exaggerated, it is a manifestation of multicomponent, multi-target, and multichannel therapy and coincides with polypharmacology along with traditional medicine. For our health, it is necessary for us to carry out more clinical trials and comprehensive pharmacological and biochemical studies to validate the clinical efficacy and the active compound(s). In summary, the results of this integrative analysis provided a comprehensive understanding of the physicochemical property and the likelihood for a natural drug, and clarified the polypharmacological mechanisms via the relationship among active ingredients, targets, and diseases. Nevertheless, the principal limitation of this analysis was the lack of validation of experiments. Conversely, it provided better guidance for subsequent experiments and also provided some novel mechanisms for clinical practice and potential active ingredients for drug discovery. Noteworthily, the reliable systems pharmacology platform of folk medicine should be constructed to develop functional foods or improve human health.

## DATA AVAILABILITY STATEMENT

All available datasets generated of this study are included in the **Supplementary Materials**.

## AUTHOR CONTRIBUTIONS

PJ, LZ, and WX conceived and proposed this work. WX, ZW, and ZX designed this research, and ZX, WZ, HC, and YG collected the data. WX, WZ, and XJ analyzed and checked the data. WX, ZW, and ZX wrote and revised the paper.

## FINDING

This work is supported by grants from National Natural Science Foundation of China (81960863, 81960870, 81760868), the Clinical Trial for the treatment of Rheumatoid Arthritis with Warming yang and Smoothening Meridians (201507001-07, registration number: ChiCTR-INR-16010290), the Construction Project of Yunnan Provincial Fund for Medical Research Center (202102AA310006), the Construction Project of National Traditional Chinese Medicine Clinical Research Base (2018 No. 131), the “Chinese Medicine Modernization Research” key project of China’s National Key R&D Programmes (2017YFC1704005), the Funding of Yunnan Applied Basic Research Projects-Union Foundation (2019FF002(−082), 2019FF002(−031)), Construction Project Funding of University Key Laboratories in Yunnan Province (2018TGZ01), and the Funding of Yunnan Provincial Health Science and Technology Plan (2018NS0045, 2018NS0046).

